# Altered functional connectivity of cortical networks in semantic variant Primary Progressive Aphasia

**DOI:** 10.1101/2020.04.13.039909

**Authors:** Haroon Popal, Megan Quimby, Daisy Hochberg, Bradford C. Dickerson, Jessica A. Collins

**Author notes:** Address correspondence to: Dr. Jessica A. Collins, MGH Frontotemporal Disorders Unit, 149 13^th^ Street, Charlestown, MA, 02129.

## Abstract

As their illness progresses, patients with the semantic variant of Primary Progressive Aphasia (svPPA) frequently exhibit peculiar behaviors indicative of altered visual attention or an increased interest in artistic endeavors. In the present study, we examined changes within and between large-scale functional brain networks that may explain this altered visual behavior. We first examined the connectivity of the visual association network, the dorsal attention network, and the default mode network in healthy young adults (n=89) to understand the typical architecture of these networks in the healthy brain. We then compared the large-scale functional connectivity of these networks in a group of svPPA patients (n=12) to a group of age-matched cognitively normal controls (n=30). Our results showed that the between-network connectivity of the dorsal attention and visual association networks was elevated in svPPA patients relative to controls. We further showed that this heightened between-network connectivity was associated with a decrease in the within-network connectivity of the default mode network, possibly due to progressive degeneration of the anterior temporal lobes in svPPA. These results suggest that focal neurodegeneration can lead to the reorganization of large-scale cognitive networks beyond the primarily affected network(s), possibly contributing to cognitive or behavioral changes that are commonly present as part of the clinical phenotype of svPPA.

## Introduction

The core symptoms of the semantic variant of Primary Progressive Aphasia (svPPA) include anomia and impaired single word comprehension (Gorno-Tempini et al., 2011; Hodges et al., 1992; Hodges & Patterson, 2007; Snowden et al., 1989; Warrington, 1975). The prominent asymmetrical atrophy and hypometabolism of the anterior temporal lobes in svPPA is usually associated with Frontotemporal Lobar Degeneration (FTLD) TDP43 neurodegenerative pathology (Josephs et al., 2011; Mesulam et al., 2014)). From a relatively early stage as the disease progresses, atrophy can also be seen in brain regions that are functionally connected with the temporal poles, including—but not limited to—the frontoinsula, subgenual anterior cingulate cortex, inferior parietal lobule, ventrolateral prefrontal cortex, and the posterior lateral and ventral temporal cortex (Brambati et al., 2009; Collins et al., 2017; Rohrer et al., 2009; Seeley, Crawford, Zhou, Miller, & Greicius, 2009).

The nomenclature used to describe the network of brain regions primarily affected by svPPA varies, with some authors highlighting overlap with anterior components of the default mode network (Bejanin et al., 2018; Collins et al., 2017; La Joie et al., 2014; Ranganath & Ritchey, 2012), or the anterior temporal/paralimbic network (Seeley et al., 2009). Although the localization of functions that support semantic memory is a matter of debate, both the default mode network (Binder et al., 2009; Jackson et al., 2016; Shapira-Lichter et al., 2013; Wirth et al., 2011) and the anterior temporal/paralimbic network (Damasio et al., 2004; Patterson et al., 2007; Visser et al., 2010) appear critically important. The temporal poles, the primary site of atrophy in svPPA (Collins et al., 2017), act as convergence zones in the brain (Tranel et al., 1998), with connectivity extending to a number of networks including the language network, visual association network, default mode network, and an anterior temporal/paralimbic network (Guo et al., 2013; Pascual et al., 2013). Thus, the neurodegenerative lesion in svPPA may compromise activity in not only the temporal poles, but potentially key nodes of the default mode, language, and visual association networks.

In addition to their well-studied language impairments, svPPA patients also frequently exhibit frank changes in the way they allocate their visual attention (Green & Patterson, 2009; Miller et al., 2000; Viskontas et al., 2011). In many patients, this alteration in visual attention takes the form of intensified focus on specific features of visual stimuli in their environment (Miller et al., 2000). For instance, the husband of an svPPA patient seen in our clinic reported that his wife devotes hours each day to compulsively watching and crushing gnats on her windowsill, possibly being driven by the visual contrast of the black insects against a white backdrop. In other patients, this change in visual attention can lead to an increased interest in visual expression; multiple anecdotes from our and others’ observations describe patients taking up an artistic hobby or dedicating more time to previous artistic pursuits (Miller et al., 2000) [https://youtu.be/3r621iKhATc]. This has been detailed in a case report of an svPPA patient (patient AA) who exhibited an interest in painting before any language symptoms or atrophy in the left anterior temporal lobe was apparent (Seeley et al., 2008). The patient had not painted for over 30 years before suddenly picking up the hobby again, which soon became an obsession.

Some of these behaviors may arise from alterations in two brain networks that normally work in concert to guide visual attention and perception in the natural environment. The visual association network, which extends along the ventral temporal lobe from early visual cortex to the temporal pole, facilitates higher-order object processing (Ungerleider & Haxby, 1994). The anterior temporal node of the visual association network, located within perirhinal cortex, is atrophied early in svPPA (Galton et al., 2001), leading to widespread functional degeneration throughout the network (Agosta et al., 2014; Guo et al., 2013). In svPPA, the functional fractionation of the visual association network may be responsible for the disproportionate impairments patients experience with object naming (Guo et al., 2013) and visual attribute reporting (Hoffman et al., 2012; Tyler & Moss, 1998). The dorsal attention network, consisting of the superior parietal lobule/intraparietal sulcus, frontal eye fields, and area MT, subserves the allocation of visual attention in exteroceptive space (Corbetta et al., 2008), and is largely spared from atrophy in svPPA. Notably, preserved grey matter density in the superior parietal lobule has been associated with faster visual search performance in svPPA patients relative to controls (Viskontas et al., 2011). In addition, svPPA patient AA had increased volume and blood flow in the right superior parietal lobule compared to a group of healthy controls, which the authors suggest may have contributed to her increased interest in visual endeavors (Seeley et al., 2008).

A defining feature of the dorsal attention network is that it activates during effortful tasks requiring attention to the external environment (Sestieri et al., 2010). In many tasks, this activation coincides with the deactivation of the default mode network (Binder et al., 1999; Corbetta & Shulman, 2002; McKiernan et al., 2003, 2006; Wirth et al., 2011). At rest, blood oxygen level dependent signal activity within these two networks appears to have inversely correlated or “anti-correlated” oscillation profiles (Fox et al., 2005; Fransson, 2005; Spreng et al., 2016). An exception to this rule is during the perception and interpretation of conceptually meaningful stimuli, during which the default mode network becomes engaged (Binder et al., 2009), and may facilitate the linking of conceptual knowledge to object representations via short range connectivity between the anterior temporal nodes of the default mode and visual association networks (Ranganath & Ritchey, 2012). In contrast, the dorsal attention network is activated during the spatial orienting response to distinctive cues, regardless of their conceptual meaning or behavioral relevance (Corbetta et al., 2008). One possibility is that the visual association network provides a sculpting influence on the reciprocal balance between the dorsal attention and default mode networks, supporting the guidance of visual attention to conceptually meaningful objects in the environment. As a corollary, we hypothesize that when connectivity between the visual association and default mode networks is compromised by a neurodegenerative lesion to the anterior temporal lobes, the dorsal attention network may be “released” from its reciprocal balance with the default mode network leading to increased connectivity with the visual association network. If this reorganization of visual-attentional networks due to neurodegeneration (RVANN) hypothesis is true, such altered brain dynamics could lead to attentional capture by perceptually distinctive features of the visual environment, regardless of their conceptual meaning.

The goal of this study was to test this RVANN hypothesis using intrinsic connectivity measured with functional MRI (fMRI). We predicted the following: 1) In healthy young adults weak between-network connectivity would be observed a) between the perirhinal node of the visual association network and the anterior temporal node of the default mode network, and b) between the posterior nodes of the visual association network and nodes of the dorsal attention network; 2) In svPPA patients, nodes of the dorsal attention network would have increased connectivity to nodes of the visual association network, compared to older controls, and 3) the increased connectivity between dorsal attention and visual association network nodes in svPPA would be associated with a functional fractionation of the anterior temporal node of the default mode network from the rest of the default mode network. This hypothesis is depicted visually in Figure 1.

**Figure 1:**
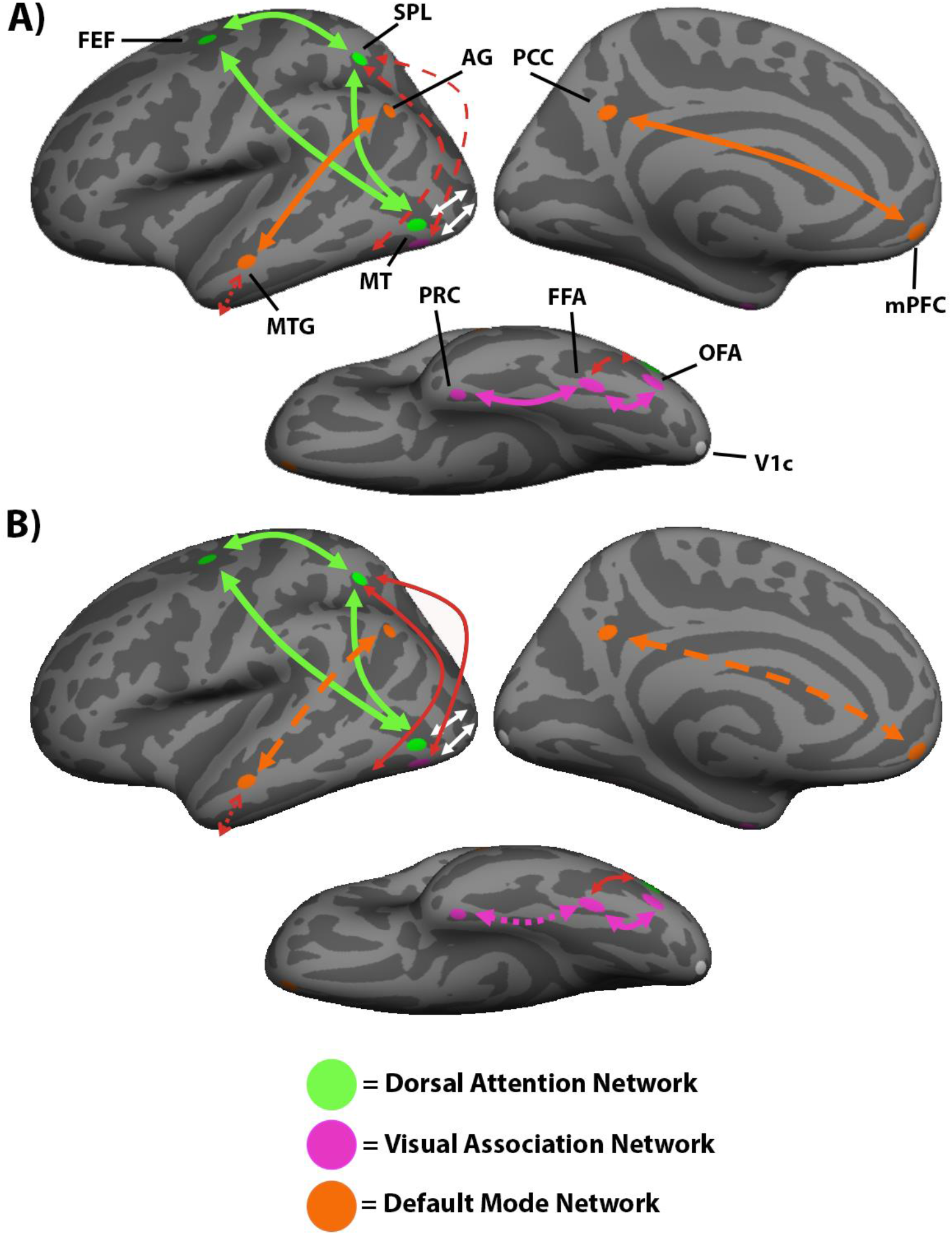
Hypothesized RVANN functional network change in svPPA patients. A) Dorsal attention network regions of interest are shown in green, visual association network regions of interest are shown in purple, and default mode network regions of interest are shown in orange. The V1c region of interest is shown in white. Arrows represent connectivity between regions of interest. In healthy adults, the dorsal attention network (green arrows) and the visual association network (purple arrows) receive input from the early visual cortex (white arrows). These networks and the default mode network (orange arrows) have strong within-network connectivity. There is normally weak connectivity between the visual association and the dorsal attention networks and between the visual association and default mode networks (shown as dashed red arrows). B) In svPPA patients, we hypothesize that the default mode network will be fractionated, due to atrophy in the lateral anterior temporal lobe. We further hypothesize that the fractionation of the default mode network will disrupt its reciprocal balance with the dorsal attention network, resulting in increased connectivity between the dorsal attention and visual association networks in svPPA (shown as solid red arrows). FEF=frontal eye fields; SPL=superior parietal lobule; MT=motion-selective cortex; FFA=fusiform face area; OFA=occipital face area; PRC=perirhinal cortex; PCC=posterior cingulate cortex; MTG=middle temporal gyrus; mPFC=medial prefrontal cortex; AG=angular gyrus; V1c=central V1.

## Materials and Methods

Three samples of participants were included in this investigation. A group of healthy younger adult controls (YC) served to test our first hypothesis regarding intrinsic connectivity within and between the dorsal attention, visual association, and default mode networks. A group of patients with a diagnosis of svPPA and a group of healthy older controls (OC), age-matched to the svPPA sample, were also included. A whole-brain group general linear model (GLM) comparing cortical thickness in the svPPA patient group to the OC group was conducted first to establish the pattern of atrophy in our patient sample. To test our second and third hypotheses, the within-network and between-network connectivity of the dorsal attention, visual association, and default mode networks were compared between the svPPA and OC groups.

### Participants

The YC group included 89 young adults (mean age = 22.4, SD = 3.34) between 18 and 33 years old, (45 female). The OC group included 30 older adults (mean age = 67.83, SD = 6.04) between 60 and 81 years old (16 females). All controls in the YC and OC groups were right-handed, native English speakers with normal vision, and had no history of neurological or psychiatric disorders.

The patient sample included 12 svPPA patients (mean age = 64.13, SD = 7.86) between 53 and 78 years old (5 females). Patients were evaluated using a structured clinical assessment performed by a behavioral neurologist and speech pathologist, as previously described (Sapolsky et al., 2010). All svPPA patients met Gorno-Tempini et al. 2011 consensus diagnostic criteria for imaging-supported svPPA based on visual inspection of a clinical MRI showing predominant left anterior temporal lobar atrophy. Ratings on the Progressive Aphasia Severity Scale (PASS) from an evaluation of each patient in the svPPA sample (Daisy Sapolsky et al., 2014) by a licensed speech language pathologist within 6 months of the patient’s MRI, along with age, sex, and years of education, is presented in Table 1. All participants gave written informed consent in accordance with the institutional review board of the Massachusetts General Hospital and Partners Healthcare System Human Research Committee.

**Table 1:** Demographic information and PASS scores for individual svPPA cases. Ed = years of education, Artic = articulation, S&G = syntax and grammar, WR = word retrieval and expression, Rep = repetition, AC = auditory comprehension, SWC = single word comprehension, Func = functional communication. For each domain: 0 = Normal, .5 = questionably impaired, 1 = mild impairment, 2 = moderate impairment, 3 = severe impairment.

| Age | Sex | Ed | Artic | Fluency | S&G | WR | Rep | AC | SWC | Reading | Writing | Func |
| --- | --- | --- | --- | --- | --- | --- | --- | --- | --- | --- | --- | --- |
| 79 | M | 18 | 0 | 0 | 0 | 1 | 0 | 0 | 0.5 | 1 | 1 | 0.5 |
| 62 | M | 18 | 0 | 0 | 0 | 0.5 | 0 | 0 | 0.5 | 0 | 0 | 0 |
| 53 | F | 18 | 0 | 0.5 | 0 | 0.5 | 0.5 | 0.5 | 0.5 | 0.5 | 0.5 | 0.5 |
| 70 | F | 12 | 0 | 0.5 | 0 | 2 | 0.5 | 1 | 2 | 2 | 2 | 1 |
| 71 | M | 12 | 0 | 0.5 | 0.5 | 1 | 0.5 | 1 | 1 | 1 | 1 | 0.5 |
| 53 | M | 20 | 0 | 0 | 0.5 | 1 | 0.5 | 1 | 1 | 2 | 0.5 | 1 |
| 55 | F | 16 | 0 | 0 | 0 | 1 | 0.5 | 0 | 0.5 | 1 | 0.5 | 0.5 |
| 68 | M | 18 | 0 | 0 | 0 | 0.5 | 0 | 0 | 0.5 | 0 | 0.5 | 0 |
| 68 | M | 17 | 0 | 0 | 0 | 0.5 | 0 | 0.5 | 0.5 | 0 | 0.5 | 0.5 |
| 65 | F | 16 | 0 | 0 | 0 | 1 | 0 | 1 | 1 | 3 | 1 | 1 |
| 67 | M | 19 | 2 | 2 | 2 | 2 | 2 | 1 | 1 | 0.5 | 1 | 1 |
| 59 | F | 12 | 0 | 0.5 | 0.5 | 2 | 0.5 | 1 | 1 | 1 | 1 | 1 |

### Scan Acquisition

All participants were scanned on a Siemens Magnetom Trio 3T scanner with a 12-channel phased-array head coil. The svPPA and OC samples were scanned using a T1-weighted MPRAGE sequence with the following parameters: TR = 2530ms, TE =3.48ms, flip angle = 7 degrees, number of interleaved sagittal slices = 176, matrix = 256×256, FOV = 256mm, voxel size = 1mm isotropic. The YC sample was scanned using a T1-weighted MEMPRAGE sequence with the following parameters: TR = 2200ms, TE = 1.54ms, flip angle = 7 degrees, number of interleaved sagittal slices = 144, matrix = 192×192, FOV = 230, voxel size = 1.2mm isotropic.

Whole brain fMRI data were acquired for the OC and svPPA samples using a T2*-weighted gradient-echo echo-planar sequence with the following parameters: TR = 5000ms, TE = 30ms, flip angle = 90 degrees, number of interleaved coronal slices = 55, matrix = 128×112, FOV = 256mm, voxel size = 2mm isotropic. Seventy-six TRs of fMRI data were acquired for a total scan duration of 6 minutes and 20 seconds.

Whole brain fMRI data was acquired for the YC sample using a T2*-weighted gradient-echo echo-planar sequence with the following parameters: TR = 3000ms, TE = 30ms, flip angle = 85 degrees, number of interleaved coronal slices = 47, matrix = 72×72, FOV = 216mm, voxel size = 3mm isotropic. 124 TRs of fMRI data were acquired for a total scan duration of 6 minutes and 12 seconds.

### fMRI Preprocessing

All intrinsic connectivity fMRI scans were preprocessed with the following parameters: removal of the first four functional volumes to allow for T1-equilibrium effects, slice timing correction (SPM2), head motion correction using rigid-body transformation in three translations and three rotations (FMRIB), spatial normalization to standard MNI-152 atlas space, resampling to 2mm isotropic voxels, spatial smoothing with a 6mm full width half max (FWHM) Gaussian kernel, and low-pass temporal filtering to remove frequencies > .08 Hz. Using linear regression, we removed sources of spurious variance and their temporal derivatives, including the six parameters from head-motion correction, the signal average over the whole brain, the average signal from a region located deep in white mater, and the average signal from ventricular cerebral spinal fluid. All subjects had less than 5 TRs of fMRI data with 5mm or more movement in any of the x, y, or z directions.

### Estimation of Intrinsic Connectivity

5mm spherical seeds and regions of interest (ROIs) were created for the dorsal attention network and the visual association network centered on MNI coordinates that have previously been shown to adequately represent their respective networks (Table 2). Seed regions for the dorsal attention network were created in the left and right superior parietal lobules (lSPL/rSPL). Other dorsal attention network ROIs included the left and right frontal eye fields (lFEF/rFEF), and left and right motion-selective cortex (lMT/rMT). Seeds for the visual association network were created in the left and right fusiform face area (lFFA/rFFA), and other visual association network ROIs included the left and right occipital face area (lOFA/rOFA) and left and right perirhinal cortex (lPRC/rPRC). Seeds for the default mode network were created in the left and right posterior cingulate cortex (lPCC/rPCC). Additional default mode network ROIs were created in the left and right medial prefrontal cortex (lmPFC/rmPFC), left and right angular gyrus (lAG/rAG), and left and right middle temporal gyrus (lMTG/rMTG, on its dorsal bank within the superior temporal sulcus) by creating 5mm spheres around the voxel showing the highest PCC connectivity in the YC group within Harvard/Oxford masks of the frontal medial cortex, angular gyrus, or the anterior and poster middle temporal gyrus, respectively. In addition, central V1 ROIs (lV1c/rV1c) were created in each hemisphere using coordinates from a previous publication (see Table 2). Correlation maps for each seed region were created by computing the Pearson’s product moment correlation ‘r’ between every voxel in the brain and the average mean signal time course of the voxels within this sphere. Positive values in this map were then converted to z-scores using Fisher’s r-to-z transformation. Maps were projected from MNI-152 space to the pial surface of the FreeSurfer average brain (fsaverage) for each ROIs’ respective hemisphere, and spatially smoothed with a 10mm FWHM kernel. For each group of subjects, maps were then thresholded to best match results from previous findings, where these ROIs were shown to be part of their discrete network (Amunts et al., 2000; Fischl et al., 2008; Fransson, 2005; Vincent et al., 2008; Wang et al., 2016). The focus of these analyses was on within-hemisphere connectivity between the ROIs, and as such, results will be reported respective to each seed’s or ROI’s hemisphere.

**Table 2:**
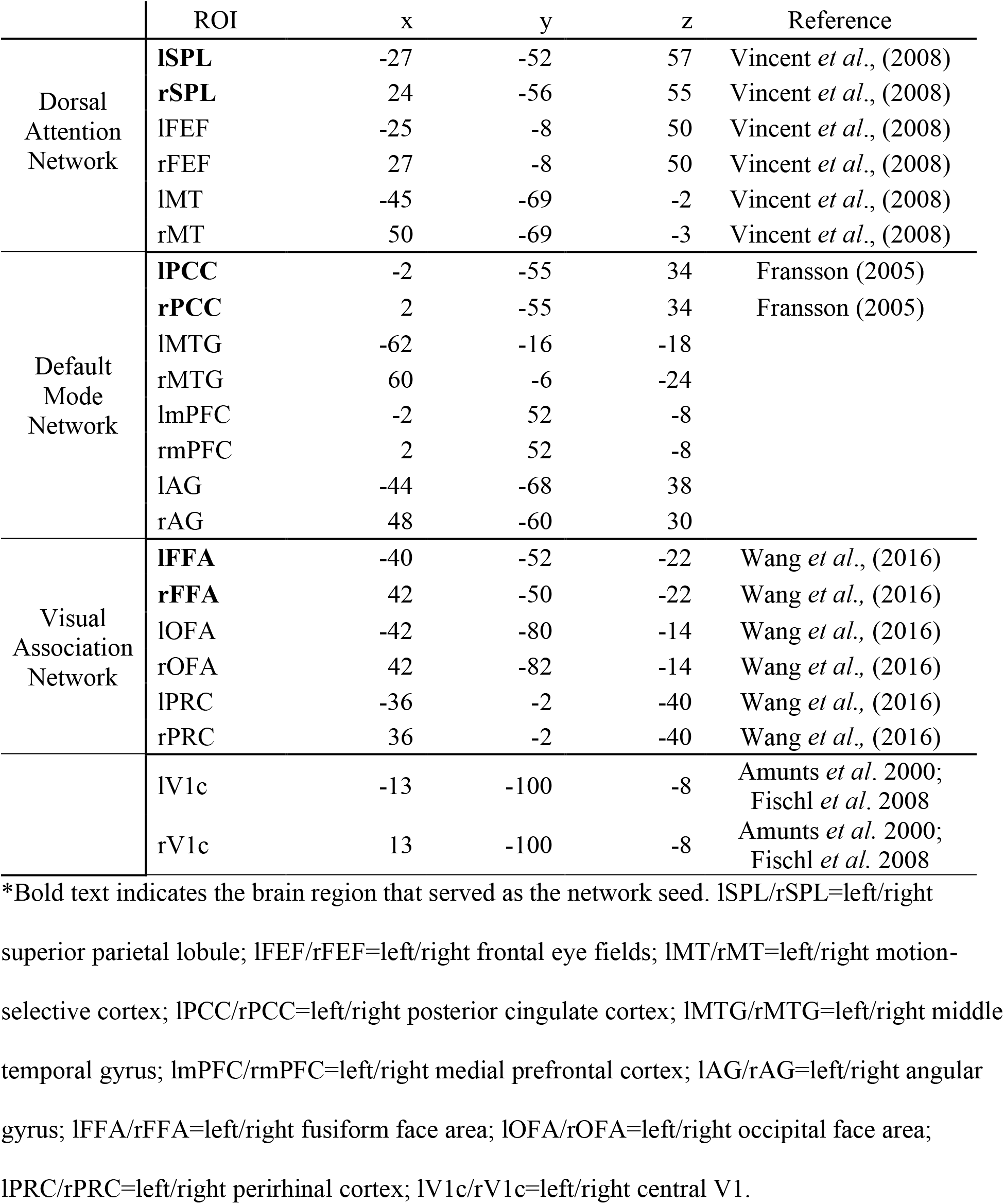
Regions of interest for the dorsal attention, visual association, and default mode networks.

In addition to the whole-brain connectivity maps, node-to-node connectivity was analyzed within each group using one-sample t-tests to assess whether between-node connectivity differed significantly from zero. The t-tests and related statistical analyses were performed using MATLAB 2018b. Two-sample t-tests were used to test hypotheses about the differences in node-to-node connectivity between the svPPA and OC groups. The z-score normalized correlations of the BOLD signal were used in all node-to-node analyses. Tests which were done to directly test our hypotheses were one-tailed t-tests, with alpha-level of *p* = .05. For post-hoc tests, two-tailed t-tests were used and to correct for multiple comparisons, a Bonferroni correction was used to determine the significance of both one-sample and two-sample t-tests, with an alpha-level of *p* = .05/55 = .00091. Tests which were not related to a priori hypotheses will be indicated as using a Bonferroni correction, with the corrected alpha-level.

### Cortical thickness analysis

The FreeSurfer analysis suite (https://surfer.nmr.mgh.harvard.edu/) was used to reconstruct cortical surfaces from the collected T1-weighted images (Dale et al., 1999; Fischl et al., 1999; Fischl & Dale, 2000; Rosas et al., 2002; Salat et al., 2004). Blind to any group affiliation, the cortical reconstruction for each individual was manually inspected to check for accuracy and to correct any errors in the automatic segmentation of grey/white matter boundaries. For each subject, the pial surface was identified using the grey/white matter boundary by a deformable surface algorithm (Fischl & Dale, 2000). Each subject’s reconstructed brain was then morphed and registered to an average spherical space (fsaverage), that optimally aligned gyral and sulcal features. The cortical thickness was calculated across the whole cortex in this new space, and a Gaussian kernel of 10mm full-width at half-maximum was used to smooth each subject’s cortical thickness map before any further analyses. A GLM was used via FreeSurfer, to compare the cortical thickness maps between the OC and svPPA groups.

### Data Availability

Imaging data are available upon request.

## Results

### Explicating dorsal attention, visual association, and default mode network connectivity in young controls

We first created whole-brain connectivity maps in the YC group bilaterally on the fsaverage surface using two canonical seeds in the dorsal attention and visual association networks: the SPL and FFA. At a typical threshold (min z = 0.10), the connectivity of SPL was largely restricted to the dorsal attention network (Fig. 2A), and the connectivity of FFA was largely restricted to the visual association network (Fig. 2B). However, at a lower threshold (min z = 0.04), the connectivity of these two seeds extended to regions typically considered part of the other network. At the lower threshold the SPL was functionally connected to regions of the visual association network including the OFA and the FFA (Fig. 2C) and the FFA was functionally connected to regions of the dorsal attention network including the SPL and the rFEF (Fig. 2D). The lower threshold of .04 was chosen because it produces results similar to what is seen with independent component analyses (Van Dijk et al., 2010).

**Figure 2:**
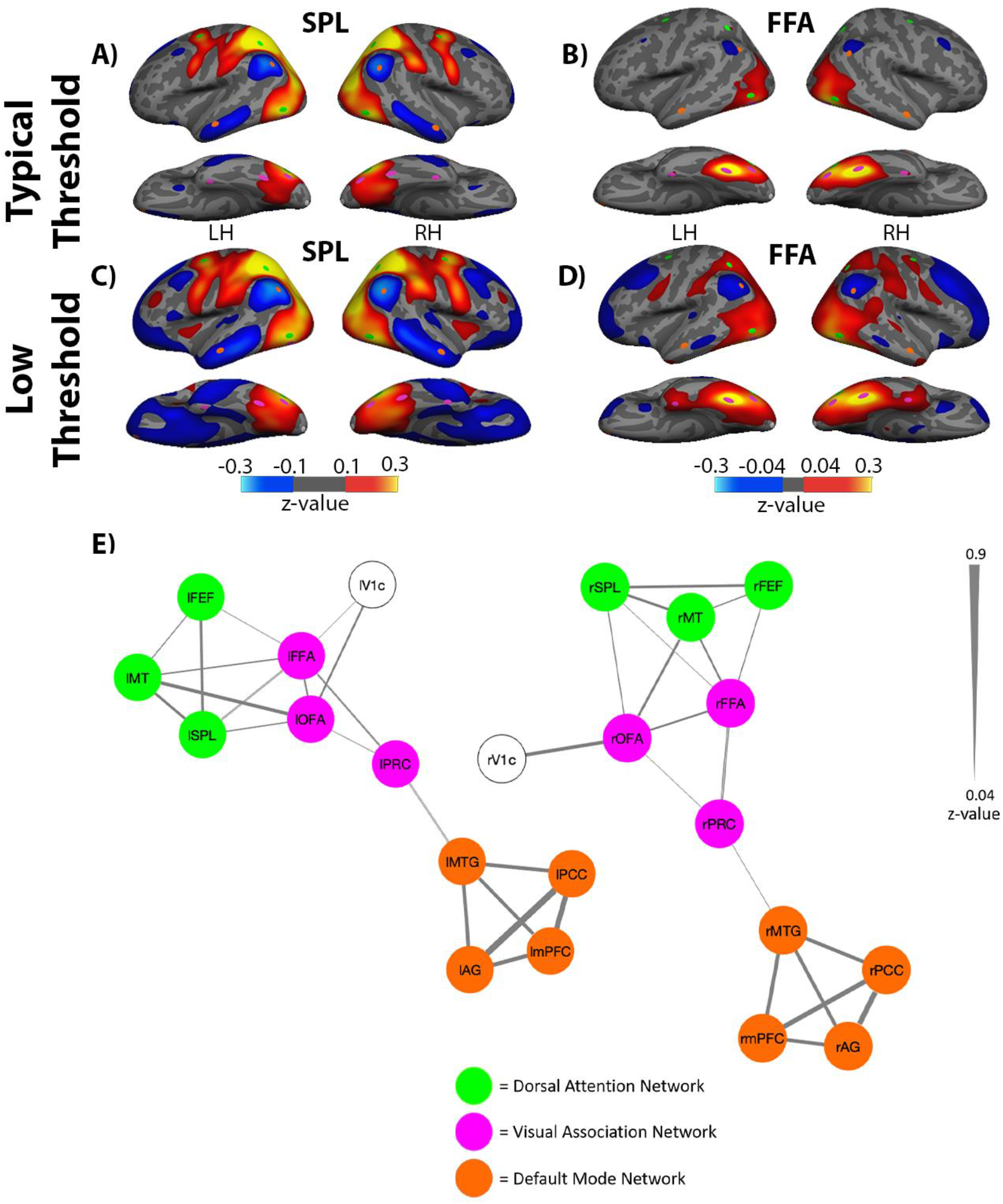
Intrinsic connectivity of dorsal attention and visual association networks in young healthy controls. A) At a typical threshold, connectivity of the SPL was mostly selective to regions within the dorsal attention network, including area MT and the FEF. At this high threshold, some with some weak between-network connectivity was observed between the SPL and OFA. The SPL also exhibited an anticorrelated pattern of connectivity with nodes of the default mode network, including AG and MTG. B) The connectivity of FFA, at a typical threshold, was mostly selective to regions within the visual association network, which included the OFA. At this high threshold, between-network connectivity was also observed between FFA and area MT. C) At a low threshold, the connectivity of the SPL extended to regions of the visual association network, including the OFA and the FFA. D) Similarly, the connectivity of the FFA extended to regions of the dorsal attention network, including bilateral SPL, and the rFEF when visualized at a lower threshold. At this lower threshold, FFA also exhibited connectivity to the anterior portions of the medial temporal lobes, including another region of the VisAN, the PRC. The FFA also exhibited an anticorrelated pattern of activity with nodes of the DMN at this lower threshold, similar to the SPL. E) Circles represent nodes for each seed or ROI, and the lines connecting the nodes represent connectivity, with thicker lines representing stronger connectivity. The dorsal attention and visual association networks showed clear within and between-network connectivity. In contrast, the default mode network was not only isolated from the other two networks, but also anti-correlated to the SPL and FFA (A-D) Weak between-network connectivity was observed between the PRC node of the visual association network and the MTG of the default mode network. Scale shows the Fischer r-z transformed connectivity values. LH=left hemisphere; RH=right hemisphere; lFEF/rFEF=left/right frontal eye frields; lSPL/rSPL=left/right superior parietal lobule; lMT/rMT=left/right motion-selective cortex; lV1c/rV1c=left/right central V1; lFFA/rFFA=left/right fusiform face area; lOFA/rOFA=left/right occipital face area; lPRC/rPRC=left/right perirhinal cortex; lPCC/rPCC=left/right posterior cingulate cortex; lmPFC/rmPFC=left/right medial prefrontal cortex; lMTG/rMTG=left/right middle temporal gyrus; lAG/rAG=left/right angular gyrus.

At a typical threshold of z = .10, the rSPL was anticorrelated with nodes of the default mode network, including the rAG and the rMTG (Fig. 2A). This replicates previous findings that demonstrate a reciprocal balance in the activity of the dorsal attention and default mode networks (Fox et al., 2005).

To further explore the within- and between-network connectivity of the dorsal attention, visual association, and default mode networks, an adjacency matrix was created by calculating the Pearson correlation between the BOLD signal time series of every node with every other node in all three networks, separately for each hemisphere. The r-values were then converted to z-scores by Fisher’s r-to-z transformation. Using the z-score adjacency matrix, a graph was created in MATLAB, using the biograph function, to visualize the connectivity between the nodes of the three networks. These graphs were thresholded so that only edges with a z-score greater than 0.04 are shown, in order to match the low threshold whole brain connectivity maps shown in Fig. 2E.

Replicating previous work, one-sample t-tests revealed within-network connectivity for all nodes of the dorsal attention, visual association, and default mode network (all *p*s < .05). Supporting our first hypothesis, between-network connectivity was observed bilaterally between area MT and the SPL within the dorsal attention network, and the FFA and OFA in the visual association network (all ps < .001; see Supplementary Table 1). Further supporting our first hypothesis, between-network connectivity was also observed bilaterally between the MTG node of the default mode network and the PRC node of the visual association network (lMTG-lPRC: *t*(88) = 5.10, *p* < .001; MTG-rPRC: *t*(88) = 2.76, *p* = .007; see Supplementary Table 1). Thus, while the FFA of the visual association network demonstrated weakly anticorrelated activity with nodes of the default mode network (Fig. 2D), the rostral temporal lobe nodes of these networks had positively correlated activity patterns, potentially representing a mechanism for the temporal pole’s function as a convergence zone. The Fischer r-z transformed connectivity values for all ROI pairs in both hemispheres is presented in Supplementary Figure 1.

### Anterior Temporal Lobe Atrophy in svPPA

A whole-brain group GLM comparing the svPPA patient group to the OC group revealed that the svPPA patients had thinner cortex in bilateral anterior temporal lobes, more prominently in the left hemisphere, as expected (Fig. 3). Atrophy in the svPPA group extended from the left temporal pole laterally to the posterior middle temporal gyrus, ventrally to the fusiform gyrus, and medially through the perirhinal cortex extending caudally to the posterior parahippocampal gyrus, as well as to the frontoinsula and orbitofrontal cortex. Less prominent atrophy was present in the right anterior temporal lobe. In both hemispheres, cortical thinning in the svPPA group included the PRC which, as discussed, is typically connected with the visual association network. Additionally, cortical atrophy in svPPA included the MTG node of the default mode network in the left and right hemispheres. That is, the two critical nodes linking the default mode network and visual association network, the PRC and the MTG, are effectively lesioned in svPPA.

**Figure 3:**
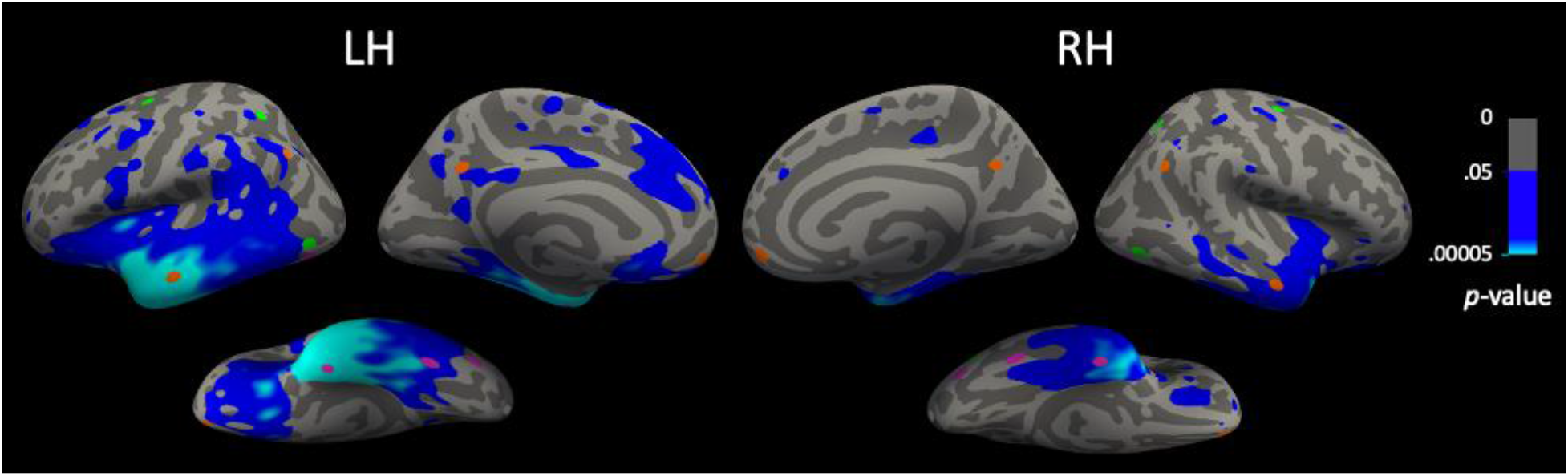
Cortical atrophy in svPPA. **A whole brain GLM** comparing cortical thickness in the svPPA group to the OC group. Areas in blue show indicate cortical thickness that was reduced (atrophic) in the svPPA group compared to OCs, thresholded at *p* < .05.

### Dorsal attention, visual association, and default mode network connectivity in svPPA and OCs

In both the svPPA and OC groups, the SPL seed showed strong connectivity with other nodes of the dorsal attention network, consistent with what was observed in the YC group (Fig. 4A-B). In the svPPA patients only, functional connectivity of the rSPL extended beyond the dorsal attention network to include ventral occipitotemporal regions rOFA and rFFA of the visual association network (Fig. 4B. The OC group did not demonstrate between-network connectivity of the rSPL and visual association network nodes (Fig. 4A). With the FFA as a seed, the OC group demonstrated similar visual association network connectivity to the YC group (Fig. 4C). However, in the svPPA group, connectivity of the rFFA extended to a number of regions of the dorsal attention network, including the rSPL and rFEF (Fig. 4D).

**Figure 4.**
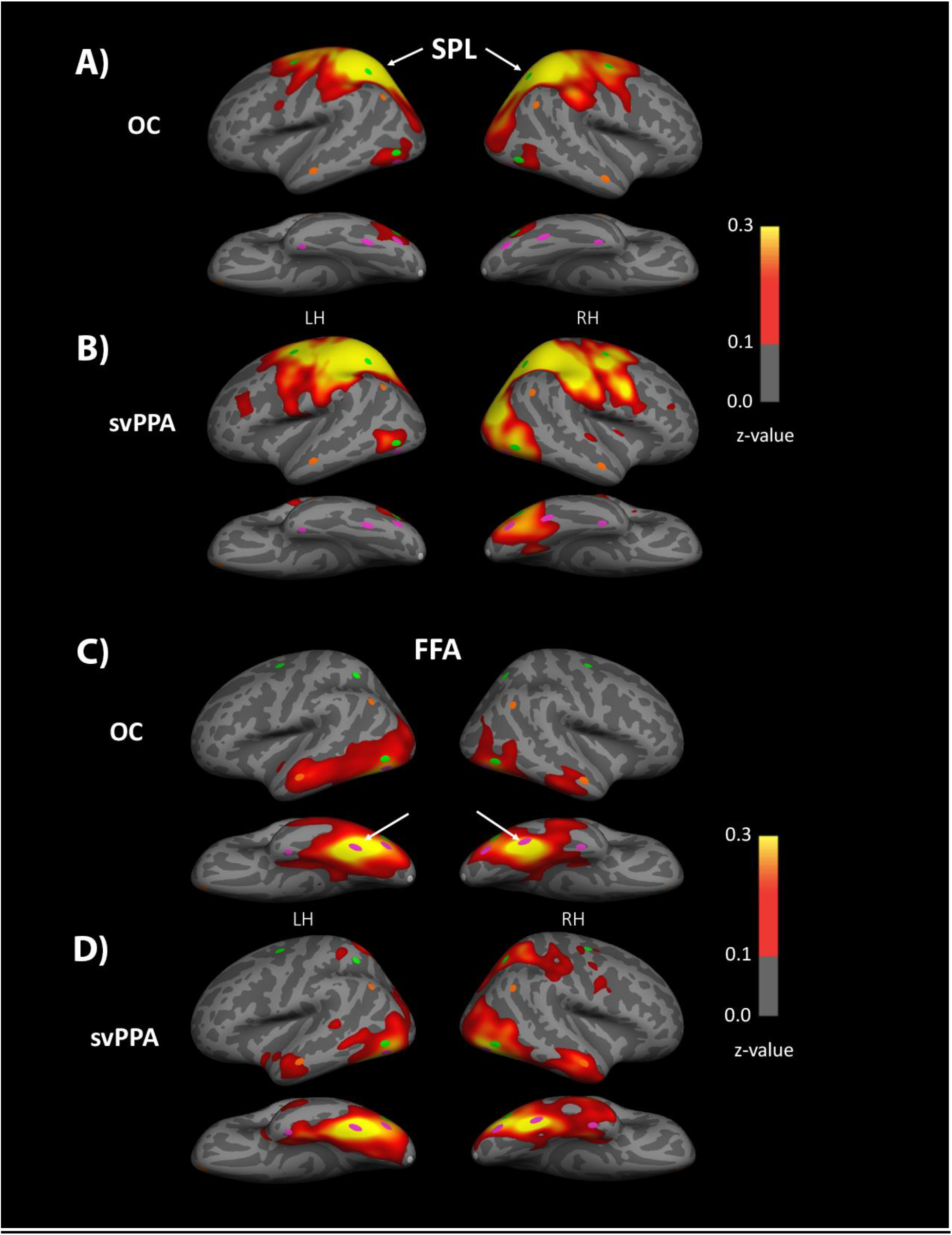
SPL and FFA intrinsic connectivity in svPPA patients and OCs. **A)** Whole-brain connectivity of the SPL in the OC group was selective to the dorsal attention network as it is in young adults, B) while connectivity of SPL in the svPPA group extended to portions of the visual association network in the right hemisphere. These maps were thresholded at z=0.10, because in YCs this threshold revealed selective functional connectivity between the SPL and other regions of the dorsal attention network. C) Whole-brain connectivity of the FFA in the OC group remained selective to the visual association network, D) while connectivity of the FFA in the svPPA group extended to portions of the dorsal attention network in the right hemisphere.

To further explore the within- and between-network dynamics of the dorsal attention, visual association, and default mode networks, adjacency matrices were created for the svPPA patient group and for the OC group displaying the connectivity of each node to every other node in all three networks. This analysis focused on nodes of the right hemisphere, as the whole-brain analysis suggested that group differences between the network architecture of the dorsal attention and visual association networks are more robust in this hemisphere. The adjacency matrices for the two groups were used to create graphs (Fig. 5) using methods identical to those used in creating the YC graphs in Fig. 2E, with the exception of thresholding so that only edges which depict Bonferroni-corrected statistically significant correlations are shown.

**Figure 5:**
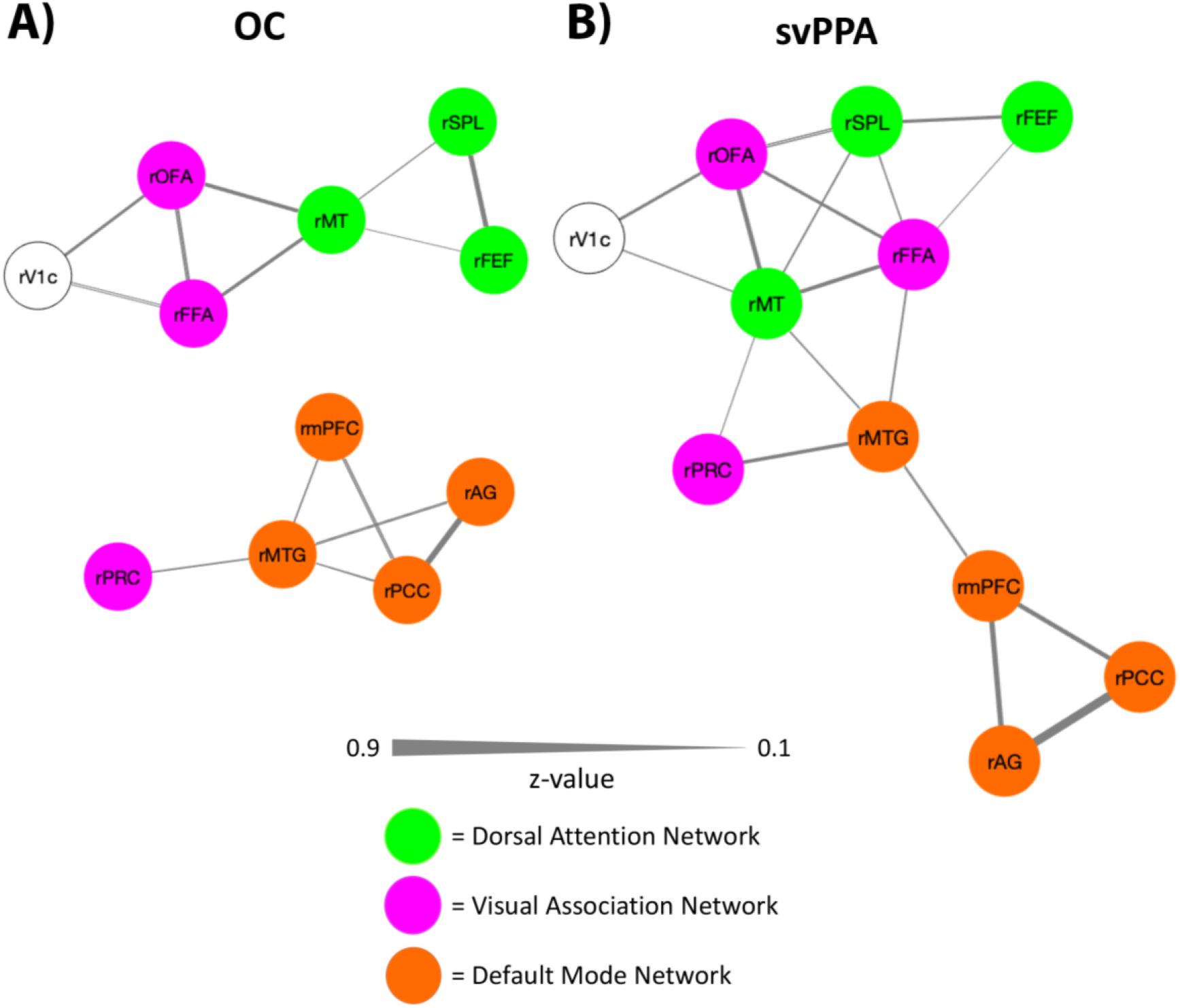
Connectivity between nodes of the dorsal attention, default mode, and visual association networks in older controls and svPPA patients. A) Older controls showed clear within-network connectivity for each network, and little between-network connectivity. B) In svPPA the within-network connectivity of the default mode network was disrupted with the rMTG node becoming fractionated from the posterior nodes of the network and demonstrating increased connectivity with visual association and dorsal attention network nodes. Between-network connectivity of the dorsal attention and visual association networks was also more prominent in the svPPA group relative to the OC group. Scale shows the Fischer r-z transformed connectivity values. rFEF=right frontal eye frields; rSPL=right superior parietal lobule; rMT=right motion-selective cortex; rV1c=right central V1; rFFA=right fusiform face area; rOFA=right occipital face area; rPRC=right perirhinal cortex; rPCC=right posterior cingulate cortex; rmPFC=right medial prefrontal cortex; rMTG=right middle temporal gyrus; rAG=right angular gyrus.

The OC group demonstrated a pattern of connectivity that was similar to the YC group, with strong within-network connectivity being exhibited by all network nodes. An exception was the rPRC, which did not demonstrate any connectivity with the rest of the visual association network nodes in the OC group, but did show connectivity to the rMTG in the default mode network, *t*(29) = 3.96, Bonferroni corrected *p* < .00091; (Supplementary Table 2). In the OC group, between-network connectivity was also observed between the rMT, in the dorsal attention networks, and two nodes of the visual association network, the rOFA, *t*(29) = 9.04, Bonferroni corrected *p* < .00091, and the rFFA, *t*(29) = 4.89, Bonferroni corrected *p* < .00091 (Supplementary Table 2; Fig. 5A).

The svPPA patient group exhibited disrupted within-network connectivity in the default mode network. Specifically, the rMTG did not retain significant connectivity with rPCC, *t*(11) = 0.67, *p* = .5177, or rAG, *t*(11) = −1.44, *p* = .1784, but retained weak connectivity to the rmPFC, *t*(11) = 1.83, *p* = .0954 (Supplementary Table 2; Fig. 5B). In the svPPA group the three other nodes of the default mode network did retain strong within-network connectivity. In the svPPA patient group, between-network connectivity was observed between the rMTG and nodes of the visual association and dorsal attention networks, including to the rPRC, *t*(11) = 3.45, *p* = .0054, rFFA, *t*(11) = 2.94, *p* = .0135, and rMT, *t*(11) = 3.27, *p* = .0074, however the results of these tests did not survive a Bonferroni correction (Supplementary Table 2).

The network graphs reinforce what is shown in Figure 4: svPPA patients showed connectivity between dorsal attention and visual association network nodes that was not present in the OC group, as was hypothesized. Specifically, in the svPPA group the rSPL showed connectivity to the rFFA, *t*(11) = 2.76, *p* = .0186, and the rFEF showed weak connectivity to the rFFA, *t*(11) = 2.02, *p* = .0686 (Supplementary Table 2). The Fischer r-z transformed connectivity values for all ROI pairs in both hemispheres is presented in Supplementary Figures 2 and 3.

### Comparison of network connectivity in svPPA patients vs OCs

To directly compare the network architecture of the svPPA and OC groups, 2-sample t-tests were conducted comparing the within- and between-network connectivity of each network node in each group. Supporting our hypothesis, svPPA patients had greater connectivity between rSPL and rFFA [t(40) = 1.72, *p* = .0463, *d* = .59] and between rFEF and rFFA [*t*(40)=2.56, *p*=.0142, *d*=0.88] relative to the OC group. Connectivity of the rSPL to the other dorsal attention network ROIs was not significantly different in the svPPA versus OC groups. Additional one-tailed t-tests supported our hypotheses that the svPPA patients have weakened connectivity between the rMTG and other default mode network nodes relative to OCs. More specifically, we showed that connectivity between the rMTG and rAG was weaker in svPPA patients compared to OCs [t(40) = 2.99, *p* = .0048, *d* = 1.02]. Connectivity between the rMTG and PCC did not differ significantly between the svPPA and OC groups [t(40) = 1.72, *p* = .0933, *d* = .59].

A whole-brain GLM was used to assess the between-group differences in the whole brain connectivity of the rSPL and the rFFA seeds for the OC and svPPA groups. A Monte-Carlo multiple comparisons correction was applied to the resulting surface maps with a vertex-wise threshold of p < .05 and a cluster-forming threshold of p < .05 (Fig. 6). For the rSPL seed, svPPA patients displayed significantly stronger connectivity to MT, OFA, and FFA relative to the OC group (cluster-wise *p*-value = .0001) (Fig. 6A). For the rFFA seed, svPPA patients demonstrated significantly stronger connectivity with the rSPL and the rFEF relative to the OC group (cluster-wise *p*-value = .0001) (Fig. 6B).

**Figure 6:**
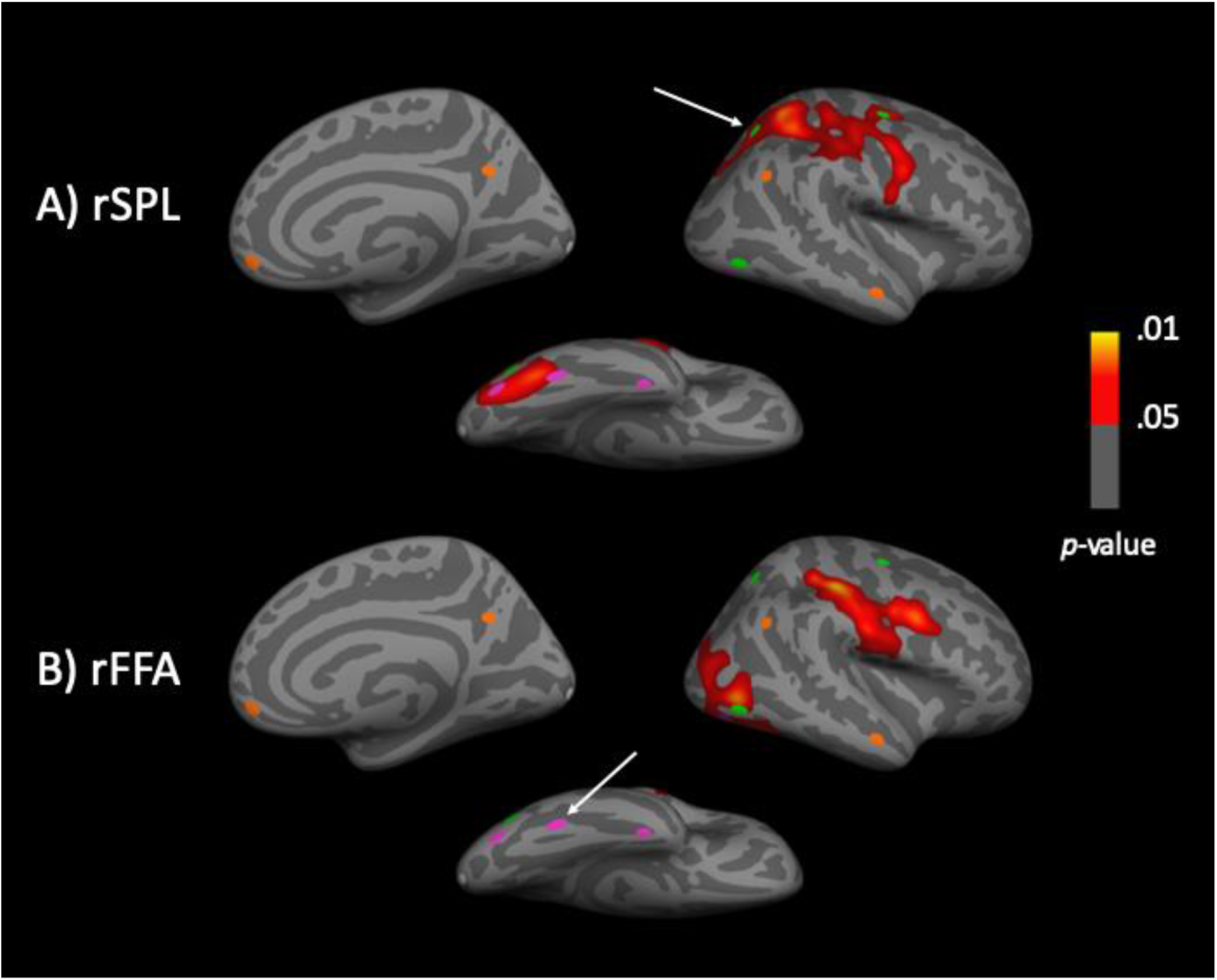
Whole-brain comparison of rFFA and rSPL connectivity in svPPA patients versus OCs. Cluster corrected whole-brain GLM analyses identified functional connectivity differences between the dorsal attention and the visual association networks in the svPPA patient versus OC groups. A) The rSPL showed greater functional connectivity with visual association network regions in the svPPA patient group compared to the OC group. B) The rFFA showed greater functional connectivity with dorsal attention network regions in the svPPA patient group compared to the OC group. Uncorrected significance maps from the original GLM are displayed for the right hemisphere, masked by significant clusters that survived the FWE correction. Arrows denote the seed region used in each analysis.

## Discussion

The goal of the present study was to investigate our RVANN hypothesis, which proposes that, in svPPA, a neurodegenerative lesion to the anterior temporal lobes leads to a functional reorganization of the brain networks supporting visual attention. We first demonstrated novel relationships between three brain networks in young healthy adults: the dorsal attention network, the visual association network, and the default mode network. Supporting our hypotheses, weak between-network connectivity was observed between the visual association and dorsal attention networks in the healthy brain. The PRC, the rostral temporal lobe node weakly connected with the visual association network, is also weakly connected to the lateral anterior temporal node of the default mode network. Supporting our RVANN hypothesis, we showed that—in patients with svPPA—the normal weak connectivity between the visual association and dorsal attention networks is aberrantly heightened, and that this increased connectivity between the visual association and dorsal attention networks in svPPA occurs in the context of a fractionated default mode network. We discuss each of these findings in greater detail below.

### Complex intrinsic relationships between dorsal attention, visual association, and default mode networks in the healthy brain

The visual association and dorsal attention networks—usually considered as distinct, separable networks—actually exhibit replicable weak connectivity with each other (Fig 2). Specifically, the SPL, a key node of the dorsal attention network, shows weak connectivity to the FFA, a key node of the visual association network, indicating that there is between-network connectivity in the healthy brain. Previous work has implicated functional interactions between the visual association and dorsal attention networks in the utilization of object-related information to guide motor actions, such as the use of a specific grasp to pick up an object (Milner, 2017; van Polanen & Davare, 2015). This evidence of between-network functional connectivity between the dorsal attention and visual association networks is consistent with evidence for structural connectivity linking the two networks (tract tracing work in non-human primates: Felleman & Van Essen, 1991; Ungerleider, Galkin, Desimone, & Gattass, 2008; Wernicke, 1881; diffusion weighted imaging in humans: Takemura et al., 2016; Yeatman et al., 2014). Connectivity between the FFA and SPL may be mediated by the vertical occipital fasciculus (VOF), which has projections to the posterior intraparietal sulcus of the dorsal attention network and the fusiform gyrus of the visual association network (Budisavljevic et al., 2018; Takemura et al., 2016; Yeatman et al., 2014). Supporting the functional evidence of dorsal attention and visual association network communication, the VOF has been implicated in reading (Greenblatt, 1973, 1976), depth perception (Oishi et al., 2018), and object related motor actions (Budisavljevic et al., 2018). Although the VOF connects regions of the visual association and dorsal attention networks, the white matter architecture that would support direct connectivity between the nodes of interest in this study (the SPL and FFA) is not understood.

The results presented here also replicated previous findings that the dorsal attention network demonstrates an anticorrelated activity pattern with the default mode network (Fig. 2). The default mode network is thought to support processes involved in semantic memory, episodic memory, prospective memory, self-referential thinking, and theory of mind (Buckner et al., 2008). When a task requiring the external orienting of attention is performed, the default mode network is deactivated and the dorsal attention network is activated (Buckner et al., 2008; Corbetta & Shulman, 2002). These task-induced activations of the dorsal attention network and deactivations of the default mode network, along with the anticorrelated intrinsic activity patterns of the dorsal attention and default mode networks, point to a reciprocal balance between these two networks in the healthy brain (Fox et al., 2005; Fransson, 2005). This reciprocal balance between the dorsal attention and default mode networks has been proposed to support the toggling of attention from internal cognitive processes to external stimuli and vice-versa (Vincent et al., 2008).

Although we also demonstrated that the posterior nodes of the visual association network are anticorrelated with the default mode network (Fig 2), we showed that the anterior ventromedial temporal node of the visual association network, the PRC, is functionally connected to the anterior lateral temporal node of the default mode network, the anterior MTG (Fig 2E). We reported this before (Pascual et al., 2013) but refined our understanding of this relationship in the context of the networks under consideration here. That is, the caudal visual association occipitotemporal cortical nodes are weakly linked to the dorsal attention network and anticorrelated with the default mode network, while the rostral ventral temporal extension of the visual association cortex is weakly connected to the default mode network. The PRC is a key node of the anterior default mode network and plays an important role in memory, especially familiarity-based item recognition memory (Collins & Dickerson, 2019; Dickerson & Eichenbaum, 2010; La Joie et al., 2014; Ranganath & Ritchey, 2012; Suzuki & Naya, 2014). The PRC has also been postulated to be the apex of the visual association network (Collins & Olson, 2014), responsible for linking complex visual representations to their associated semantic features (Barbeau et al., 2008; Martin et al., 2013; Taylor et al., 2006). This process in humans may be mediated by short-range connectivity of PRC with adjacent regions in the lateral anterior temporal lobe, including MTG, integrating high-level object perception with semantic and episodic memory (Miyashita, 2019). Supporting this notion, tract tracing work in macaques has established the existence of direct structural connections from the PRC to TGvg and TGsts (Kondo et al., 2003, 2005; Saleem et al., 2008). Notably, the anterior MTG and the PRC both succumb to neurodegeneration early in the disease course of svPPA (Collins et al., 2017; Davies et al., 2005; Mummery et al., 2000), providing a potential mechanism for these two regions to become fractionated from each other, and from their respective networks.

### Neurodegeneration of anterior temporal hubs disrupts within- and between-network connectivity in svPPA

Consistent with our hypotheses, in svPPA, the normally weak connectivity between nodes of the dorsal attention and visual association networks is heightened relative to healthy older adults (Figs. 4-6). Specifically, we showed that—in svPPA patients—the rSPL had heightened connectivity with rFFA and rOFA, and the rFEF had heightened connectivity with rFFA. Our results expand upon previous findings suggesting that the SPL might play a critical role in driving altered visual behavior in svPPA. In a previous case study of svPPA, increased blood flow and volume in the rSPL was associated with increased artistic abilities (Seeley et al., 2008). In addition, in a group of patients with svPPA, Viskontas *et al*. (2011) has shown that the volume of the right and left SPL predicts performance on a conjunction search task, in which participants were measured on how quickly they could find target letter stimuli of a particular color (e.g. green Ns among brown Ns and green Xs). The SPL is a heteromodal association area that has been shown to play a critical role in visual search and the allocation of visual attention in healthy adults (Corbetta et al., 2002; Downar et al., 2000; Kim et al., 1999); we speculate that aberrant neurodegeneration-related increases in the SPL’s connectivity with nodes of the visual association network may lead to attentional capture by simple perceptually rich stimuli, without regard to their meaning.

We also showed that the within-network connectivity of the default mode network becomes disrupted in svPPA, with the rMTG becoming functionally lesioned from the more posterior nodes of the default mode network including the PCC and AG. These results are consistent with previous work showing a selective vulnerability of anterior temporal default mode network regions in svPPA, in contrast to posterior and medial regions of the default mode network, which are commonly atrophied in Alzheimer’s disease and spared in svPPA (La Joie et al., 2014; Ranganath & Ritchey, 2012). The anterior MTG in the default mode network was of interest in the current study because it is consistently atrophied in svPPA (Bejanin et al., 2018; Collins et al., 2017; La Joie et al., 2014; Pascual et al., 2013; Ranganath & Ritchey, 2012). We hypothesized that when this region of the default mode network is lesioned, connectivity between the visual association and default mode networks would be compromised, and the dorsal attention network would be “released” from its inhibition by the default mode network leading to increased connectivity between the dorsal attention and visual association networks. In support of this hypothesis, previous research in healthy young adults has shown that inhibiting the anterior default mode network through repetitive transcranial magnetic stimulation to the left ATL can lead to the emergence of savant-like visual and numerosity skills. More specifically, in one study participants were able to more accurately guess the number of objects presented on a computer screen (between 50-150) in a 1-2 second period following repetitive transcranial magnetic stimulation to the left ATL (Snyder et al., 2006). In a separate study participants had an increased ability to create naturalistic drawings after repetitive transcranial magnetic stimulation to the left ATL (Snyder, 2009). While the functional network basis of these effects remains unclear, the results of these transcranial magnetic stimulation studies are consistent with the results presented here in demonstrating that a lesion to the lateral anterior temporal node of the default mode network can affect visual behavior.

### Limitations and Future directions

In svPPA patients with primarily left hemisphere anterior temporal lobe atrophy, subtle right anterior temporal atrophy can be seen in a circumscribed manner before it spreads posteriorly to cover more of the temporal lobe (Rogalski et al., 2011). One reason our results were confined to the right hemisphere may be that the left hemisphere is too atrophied to reveal these kinds of changes in functional connectivity, even though most of these patients were mildly clinically impaired. It is common for mildly clinically impaired svPPA patients to have relatively prominent left anterior temporal atrophy. If patients could be identified at an earlier stage of neurodegeneration, we would expect to see similar connectivity patterns in the left hemisphere; possibly as early as the first changes in visual attention are reported, which can sometimes be before language symptoms are present (Seeley et al., 2008).

In the present study we did not collect behavioral measures of visual attention, and thus we cannot directly relate the changes in brain networks observed in our svPPA sample to changes in behavior. In future work, we aim to study the changes in visual attentional capture in svPPA using behavioral and eye tracking measures, with the goal of better understanding how large-scale functional network changes due to neurodegeneration can give rise to alterations in complex visual behavior.

## Abbreviations

svPPA: semantic variant Primary Progressive Aphasia
OC: older controls
YC: young controls
DAN: dorsal attention network
VisAN: visual association network
DMN: default mode network

## Supplementary Materials

**Supplementary Table 1.** Between-network ROI-to-ROI connectivity statistics for young healthy controls. Degrees of freedom for all statistical tests = 88.

| Node to Node Connectivity | t-stat | p-value | Mean | 95% CI |
| --- | --- | --- | --- | --- |
| IFFA - ISPL | 5.67 | < .001 | 0.10 | [0.07, 0.14] |
| IFFA - IMT | 9.53 | < .001 | 0.20 | [0.16, 0.25] |
| rFFA - rSPL | 3.91 | < .001 | 0.07 | [0.04, 0.11] |
| rFFA - rMT | 12.13 | < .001 | 0.28 | [0.24, 0.33] |
| IOFA - ISPL | 8.24 | < .001 | 0.18 | [0.14, 0.22] |
| IOFA - IMT | 13.35 | < .001 | 0.29 | [0.25, 0.34] |
| roFA - rSPL | 7.68 | < .001 | 0.17 | [0.12, 0.21] |
| roFA - rMT | 16.36 | < .001 | 0.36 | [0.31, 0.40] |
| IMTG - IPRC | 5.10 | < .001 | 0.08 | [0.05, 0.11] |
| rMTG - rPRC | 2.76 | .007 | 0.06 | [0.02, 0.10] |

ISPL/rSPL=left/right superior parietal lobule; IMT/rMT=left/right motion-selective cortex;
IFFA/rFFA=left/right fusiform face area; IOFA/roFA=left/right occipital face area;
IPRC/rPRC=left/right perirhinal cortex; IMTG/rMTG=left/right middle temporal gyrus;
LAG/rAG=left/right angular gyrus.

**Supplementary Table 2.**
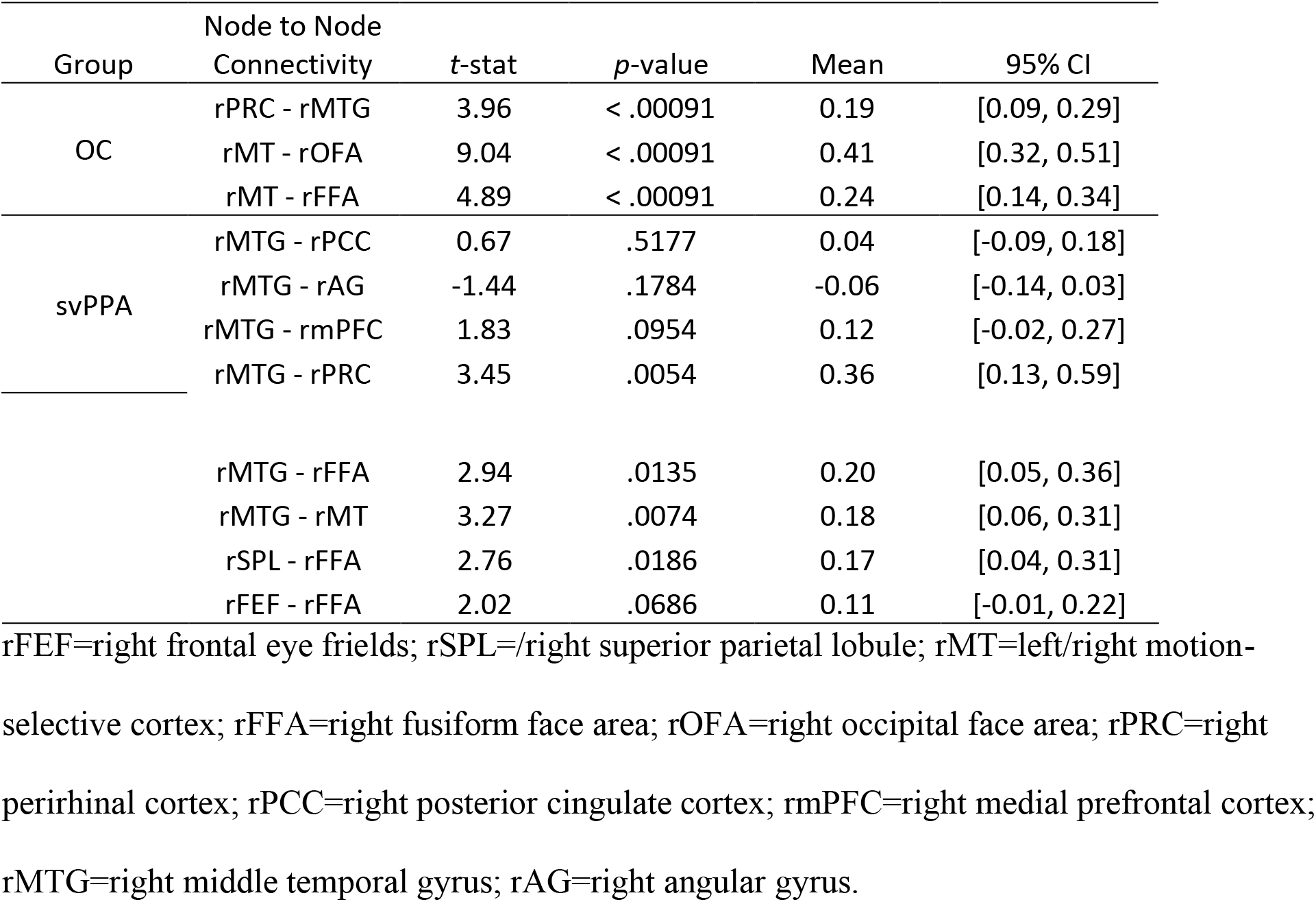
RVANN ROI-to-ROI connectivity statistics for older healthy controls and semantic variant Primary Progressive Aphasia patients. Degrees of freedom for OC statistical comparisons = 29. Degrees of freedom used for svPPA statistical comparisons = 11.

| Group | Node to Node Connectivity | t-stat | p-value | Mean | 95% CI |
| --- | --- | --- | --- | --- | --- |
| OC | rPRC - rMTG | 3.96 | < .00091 | 0.19 | [0.09, 0.29] |
|  | rMT - roFA | 9.04 | < .00091 | 0.41 | [0.32, 0.51] |
|  | rMT - rFFA | 4.89 | < .00091 | 0.24 | [0.14, 0.34] |
| svPPA | rMTG - rPCC | 0.67 | .5177 | 0.04 | [-0.09, 0.18] |
|  | rMTG - rAG | -1.44 | .1784 | -0.06 | [-0.14, 0.03] |
|  | rMTG - rmPFC | 1.83 | .0954 | 0.12 | [-0.02, 0.27] |
|  | rMTG - rPRC | 3.45 | .0054 | 0.36 | [0.13, 0.59] |

Running head: Functional connectivity in svPPA
|  |  |  |  |  |
| --- | --- | --- | --- | --- |
| rMTG - rFFA | 2.94 | .0135 | 0.20 | [0.05, 0.36] |
| rMTG - rMT | 3.27 | .0074 | 0.18 | [0.06, 0.31] |
| rSPL - rFFA | 2.76 | .0186 | 0.17 | [0.04, 0.31] |
| rFEF - rFFA | 2.02 | .0686 | 0.11 | [-0.01, 0.22] |

**Supplementary Figure 1:**
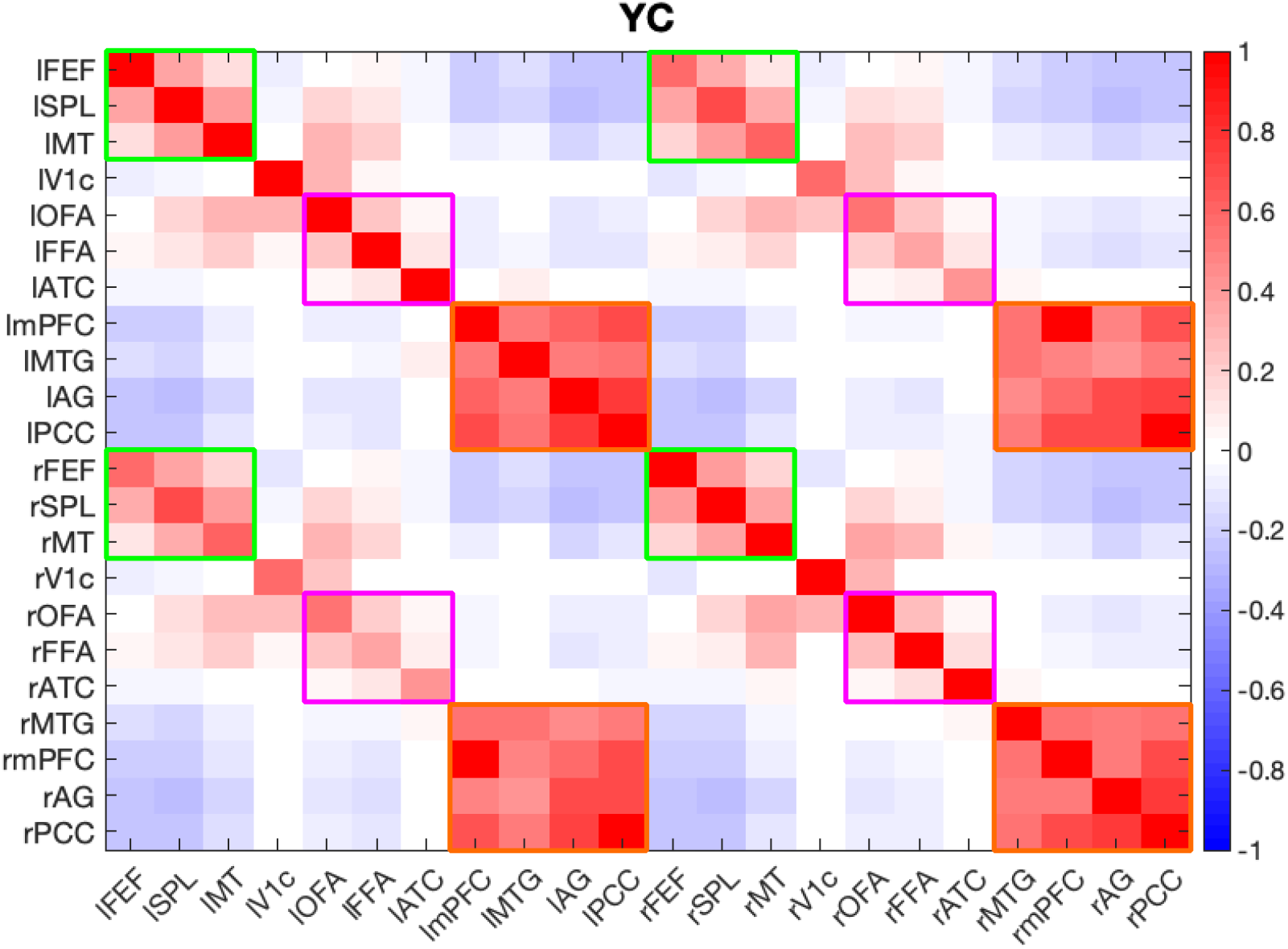
Connectivity between all nodes of the dorsal attention, default mode, and visual association networks in young controls. rFEF=right frontal eye frields; rSPL=/right superior parietal lobule; rMT=left/right motion-selective cortex; rFFA=right fusiform face area; rOFA=right occipital face area; rPRC=right perirhinal cortex; rPCC=right posterior cingulate cortex; rmPFC=right medial prefrontal cortex; rMTG=right middle temporal gyrus; rAG=right angular gyrus; lV1c/rV1c=left/right central V1.

**Supplementary Figure 2:**
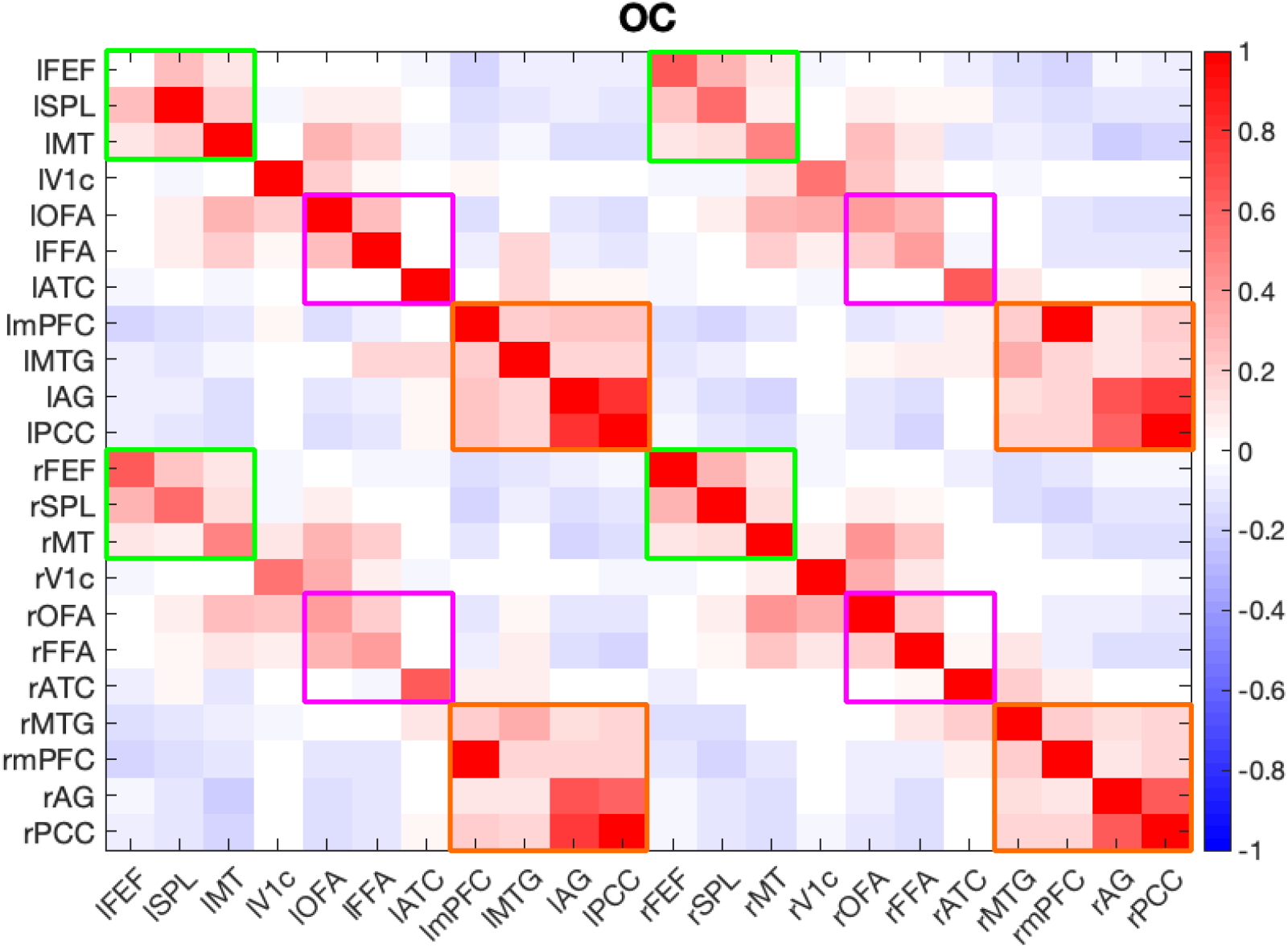
Connectivity between all nodes of the dorsal attention, default mode, and visual association networks in older controls. rFEF=right frontal eye frields; rSPL=/right superior parietal lobule; rMT=left/right motion-selective cortex; rFFA=right fusiform face area; rOFA=right occipital face area; rPRC=right perirhinal cortex; rPCC=right posterior cingulate cortex; rmPFC=right medial prefrontal cortex; rMTG=right middle temporal gyrus; rAG=right angular gyrus; lV1c/rV1c=left/right central V1.

**Supplementary Figure 3:**
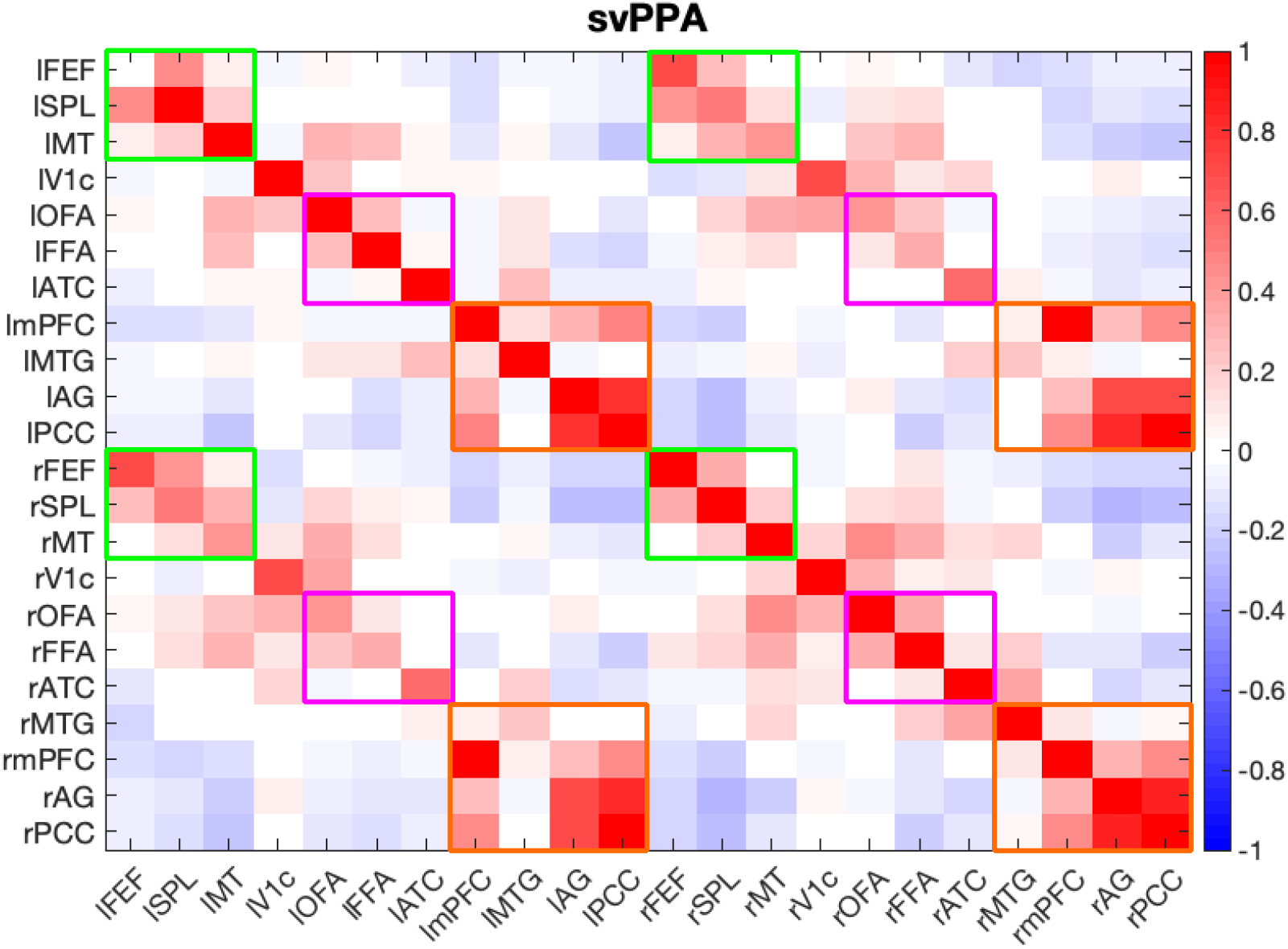
Connectivity between all nodes of the dorsal attention, default mode, and visual association networks in svPPA patients. rFEF=right frontal eye frields; rSPL=/right superior parietal lobule; rMT=left/right motion-selective cortex; rFFA=right fusiform face area; rOFA=right occipital face area; rPRC=right perirhinal cortex; rPCC=right posterior cingulate cortex; rmPFC=right medial prefrontal cortex; rMTG=right middle temporal gyrus; rAG=right angular gyrus; lV1c/rV1c=left/right central V1.

